# Transcriptional and translational profiling in yeast reveals the use of diverse genome decoding mechanisms to generate functionally distinct proteoforms

**DOI:** 10.1101/2022.03.25.485750

**Authors:** Darren A Fenton, Michał Świrski, Patrick B F O’Connor, Stephen J Kiniry, Audrey M Michel, Joanna Kufel, Pavel V Baranov, John P Morrissey

## Abstract

The coding potential of the eukaryotic genome can be greatly expanded by the regulated use of mechanisms that generate more than one protein product from a gene. We combined techniques for mapping 5’ and 3’ ends of RNA transcripts with ribosome profiling to study the organisation of protein coding gene expression in the yeast *Kluyveromyces marxianus*. We uncovered over 1000 cases of novel proteoforms due to use of alternative transcription or translation start sites, identified 800 translated upstream open reading frames, observed surprising translation of antisense RNAs, and discovered a novel case of programmed ribosomal frameshifting. In some cases, features are conserved across yeast species, whereas others are species-specific. This offers new possibilities to explore the evolution of genomes and gene regulation in budding yeasts. Our analysis also enabled us to improve the genome annotation of *K. marxianus* by adding or correcting annotations of over 300 protein coding genes. The processed data has been made available on the GWIPS-viz and Trips-Viz browsers, thus providing an accurate data-driven annotation of transcripts and their protein coding regions along with quantitative information on their transcription and translation.

## INTRODUCTION

*Kluyveromyces marxianus* is a budding yeast in the Saccharomycetaceae family. Although readily isolated from decaying plant matter, it is believed that some *K. marxianus* lineages were domesticated by early dairy farmers several thousand years ago and it is traditional associated with fermented dairy products (Ortiz-Merino *et al.*, 2018; Varela *et al.*, 2019). More recently, its capacity for rapid growth, thermotolerance and other relevant traits has garnered much attention in the biotechnology sector (Fonseca *et al.*, 2008; Lane and Morrissey, 2010; Morrissey *et al.*, 2015; Varela *et al.*, 2017; Karim, Gerliani and Aïder, 2020). This has led to substantial progress in the development of gene engineering and synthetic biology tools (Nambu-Nishida *et al.*, 2017; Cernak *et al.*, 2018; Rajkumar *et al.*, 2019; Rajkumar and Morrissey, 2022), meaning that it is now relatively straightforward to reprogramme strains as cell factories for industrial biotechnology applications (Liu and Nielsen, 2019; Rajkumar and Morrissey, 2020; Baptista, Cunha and Domingues, 2021; Patra *et al.*, 2021). Other developments at the physiological level are improving the potential for large-scale fermentation of *K. marxianus* under industrial conditions (Dekker *et al.*, 2021). There have also been several genome-wide studies that explored gene expression at the transcriptional level (reviewed in (Ha-Tran, Nguyen and Huang, 2020).

To date, 17 *K. marxianus* genomes have been published, with 3 strains (NBRC 1777, DMKU-1042 and FIM1) having fully sequenced genomes, annotated and assembled on a chromosomal level (Inokuma *et al.*, 2015; Lertwattanasakul *et al.*, 2015; Mo *et al.*, 2019). Similar to other pre-whole genome duplication yeast in the Saccharomycetaceae, *K. marxianus* possesses eight nuclear chromosomes and a haploid genome of ~10.9 Mb in size, although variations in ploidy and aneuploidy are common in the domesticated dairy strains (Fasoli *et al.*, 2015; Ortiz-Merino *et al.*, 2018). Annotation of the sequenced genomes indicates that there are ~5000 protein-coding genes in *K. marxianus*, lower than the ~6000 in *S. cerevisiae*. There have been only limited efforts to explore the entire *K. marxianus* genome to establish the basis of its unique or interesting traits. Most previous studies were narrowly focused, for example on the expansion and diversification of sugar transporters (Knoshaug *et al.*, 2015; Varela *et al.*, 2017, 2019; Donzella *et al.*, 2021). In a recent comparative omics study, we discovered that genes that were evolutionarily young and / or unique to *K. marxianus* are overrepresented among genes that display differential expression when growing under stressful conditions (Doughty *et al.*, 2020). We also established that at least one of these genes is required for competitive growth at high temperature (Montini *et al.*, 2022). These findings reinforce the need to thoroughly explore and characterise the *K. marxianus* genome as many of the unique phenotypic traits in *K. marxianus* could be due to genes not yet characterised in any species, or indeed to known genes that are regulated differently or encode proteins with alternative functionality.

Genome annotation in yeast is generally homology-based, which is rapid but brings some limitations, especially when dealing with less well-characterised species. Among the limitations of this conventional protein-coding gene annotation approach, is the difficulty in considering the diversity of proteoforms that can be encoded within a single gene locus. This is illustrated well in *Saccharomyces cerevisiae* where studies revealed a range of different types of proteoforms generated with alternative translation initiation codons (Monteuuis *et al.*, 2019), translational readthrough (Namy *et al.*, 2003), and frameshifting (Atkins *et al.*, 2016). Proteoforms with different N-termini can arise through either transcription or translation-based mechanisms and, amongst other roles, are known to have different localisation within the cell, due to a targeting signal present at the N-terminus of the longer proteoform. For example, *S. cerevisiae HTS1* uses alternative transcription start sites that give rise to long and short mRNA isoforms that are then translated from different start codons 60 nt apart (Natsoulis, Hilger and Fink, 1986). In contrast, translation of the *S. cerevisiae ALA1* mRNA involves leaky scanning, whereby a proportion of ribosomes initiate translation at upstream near-cognate start codons (ACG), while remaining ribosomes continue scanning and initiate translation downstream at the main AUG start codon (Tang *et al.*, 2004). Other translational mechanisms can result in proteoforms with alternative C-termini. This includes stop codon readthrough, whereby ribosomes fail to terminate at stop codons and continue translation, or ribosome frameshifting, whereby a ribosome repositions and translates in the −1 or +1 open reading frame (ORF) (Namy *et al.*, 2003; Atkins *et al.*, 2016). In addition to enabling synthesis of alternative proteoforms, these mechanisms could provide regulatory sensing of cellular conditions. The best-known example is the regulation of intracellular levels of polyamines via +1 frameshifting that takes place during translation of *OAZ1* mRNA (Palanimurugan *et al.*, 2004; Ivanov, Gesteland and Atkins, 2006), which is conserved from yeast to humans (Ivanov and Atkins, 2007). Other important translated regions that can be missed in gene annotation include small upstream open reading frames (uORFs) within 5’ leaders of mRNAs. In *S. cerevisiae* these uORFs have been shown to regulate translation of mRNAs including for example *CPA1*, which encodes an arginine attenuator peptide, where ribosome stalling at an uORF is regulated via intracellular arginine levels (Gaba *et al.*, 2001; Gaba, Jacobson and Sachs, 2005).

Regarding studies of gene expression, massively parallel sequencing facilitated a switch in research focus from gene-specific studies to the genome-wide scale. Transcriptomic methods such as RNA-seq can be used to measure gene expression (Wang, Gerstein and Snyder, 2009) or to capture transcript start and polyadenylation sites, revealing alternative mRNA isoforms based on 5’ or 3’ ends (Adiconis *et al.*, 2018; Yu *et al.*, 2020). Ribosome profiling is another genome-wide tool to measure gene expression. This tool captures and locates the positions of translating ribosomes, revealing precise regions of the genome that are translated at a moment in time. Using ribosome profiling (Ribo-Seq), it is possible to investigate the translational control of specific mRNAs, measure gene expression changes at a translational level, determine the translation efficiency of a transcript, and identify novel coding regions (Ingolia *et al.*, 2009; Michel and Baranov, 2013; McManus *et al.*, 2014; Brar and Weissman, 2015; Kiniry, Michel and Baranov, 2020). Ribosome profiling was first carried out in *S. cerevisiae* (Ingolia *et al.*, 2009), then later in the other yeasts, *Saccharomyces paradoxus* (McManus *et al.*, 2014), *Schizosaccharomyces pombe* (Duncan and Mata, 2014), *Saccharomyces uvarum* (Spealman *et al.*, 2018), *Komagataella phaffii* (Alva, Riera and Chartron, 2021) and *Candida albicans* (Sharma *et al.*, 2021). Using ribosome profiling, it has been possible to uncover many non-canonical aspects of gene structure and regulation. In *S. cerevisiae*, the approach revealed a wide-range of translated upstream open reading frames (uORFs) within 5’ leaders of mRNAs (Ingolia *et al.*, 2009), widespread non-AUG initiation encoding unannotated proteoforms (Monteuuis *et al.*, 2019; Eisenberg *et al.*, 2020) and identified a number of previously unknown translated small ORFs (Smith *et al.*, 2014). In *K. marxianus*, genome-wide studies of gene expression have focused mainly on RNA-seq (reviewed in (Ha-Tran, Nguyen and Huang, 2020)), ignoring these more complex mechanisms as they only reveal relative mRNA abundances.

We applied the ribosome profiling method to *K. marxianus* and established a protocol and pipeline for ribosome profiling in this yeast (Fenton et al., 2022). To gain a better understanding of the Kluyveromyces transcriptome and translatome, we employed a combination of transcriptomics techniques and ribosome profiling that allowed us to improve its genome annotation by increasing its accuracy, and inclusion of alternative proteoforms. In addition to depositing our primary data and annotations into standard depositories we generated a *K. marxianus* entry at GWIPS-viz (Michel *et al.*, 2014, 2018) and Trips-Viz (Kiniry *et al.*, 2019, 2021) browsers where processed data can be freely and conveniently explored alongside the new annotation.

## MATERIALS AND METHODS

### RNA-Seq and Ribosome Profiling

RNA-Seq and ribosome profiling was carried out as in (Fenton *et al.*, 2022). Splice junctions were identified for novel intron containing genes and genes which required splice site corrections using the splice aware STAR RNA-seq aligner (Dobin *et al.*, 2013). In the ribosome profiling analysis, RPFs that failed to align to the original CDS regions of DMKU3-1042 annotation were aligned to the reference genome. These alignments were then split into windows using Bedtools (Quinlan and Hall, 2010). Windows were ranked based on the number of alignments and the top candidates were visually assessed using a genome browser (GWIPS-viz (Michel *et al.*, 2014)) where we created a database for *K. marxianus* DMKU-1042 genome. For the BLASTP and TBLASTN heatmaps, custom databases containing annotated protein sequences and genome assemblies were created for use with BLASTP and TBLASTN, respectively (Altschul *et al.*, 1990). For BLASTP and TBLASTN, the following parameters were specified, -seg no, -threshold 11, -max_hsps 1 and -outfmt 6. An e value filter of =<0.01 was applied to blast results.

### TSS-Seq

In order to precisely characterise Transcription Start Sites, publicly available TSS-seq data originating from *K. marxianus* DMKU3-1042 was used (Lertwattanasakul *et al.*, 2015). Data were downloaded from the NCBI SRA repository, adaptor removal and quality trimming was performed with cutadapt, followed with rRNA removal and genome alignment with bowtie. While doing this analysis, we noticed that accession numbers of raw data deposited in SRA do not match expression profiles of conditions discussed in the original paper (Lertwattanasakul *et al.*, 2015) and were evidently mislabelled during sequence deposition. For our analysis, we reassigned the samples to the correct condition, detailed in Supplementary material. Resulting alignments were used for transcriptional units (Transcription Start Region -– TSR) detection by clustering reads with Bioconductor package CAGEr (Haberle *et al.*, 2015). TSRs were subsequently assigned to the nearest coding sequence (CDS) with a minimum relative expression cut-off of 0.05 and 1 TPM was applied to filter out lowly expressed TSRs or unreliable clusters (see Supplementary Figure 5).

### Identification of PAS

For identification of polyadenylation sites, we used our own polyA-enriched RNA-seq data (Fenton *et al.*, 2022). Reads that aligned to the genome were discarded as these are sequences that do not contain polyA tails. From the remaining reads, all trailing A nucleotides were trimmed from 3’ ends of reads and aligned once again to the genome revealing PAS. Aligned reads were processed analogously to the TSS-seq reads: clustering has been performed with CAGEr and followed with assignment to nearest CDS with minimum relative expression cut-off of 0.05 and 1 TPM. Remaining clusters were assumed as *bona-fide* polyadenylation sites (PAS, see Supplementary Figure 6).

### Multiomics plots

Briefly, BAM files were processed with ORFik bioconductor package to generate P-site ribo-seq profiles and RNA-seq coverage profiles (Tjeldnes *et al.*, 2021). In the process of mining the data, RiboCrypt: R package NGS data visualization tool was developed. It takes use of ORFik data management and processing and ggplot2 combined with plotly for data display. RiboCrypt GitHub repository is available at https://github.com/m-swirski/RiboCrypt. The profiles are characterised by very sharp peaks, making profiles less-readable when zoomed-out. Thus a sliding window mean of a profile was used to decrease resolution and increase clarity of a picture. To display coverage we employed the stacked method to avoid blurring the plot by area overlapping.

### Non-canonical translation detection

All possible ORFs starting from any one of the near-cognate codons (differing from AUG by one nucleotide, including AUG) in the *K. marxianus* genome and transcriptome were found with ORFik package (Tjeldnes *et al.*, 2021). Naturally, it resulted in finding multiple ORFs sharing a stop codon.

Subsequently, a P-site profile was generated for the longest ORF for each stop codon and a set of parameters was calculated for all nested ORFs. P-site score: % of in-frame reads, read count – given as reads per kilobase (RPK), in-frame coverage fold-change between a region between Nth and Nth+1 start codon and first to Nth-1 start codon. Unique P-site score: P-site score calculated for a region between Nth and Nth+1 start codon. Additionally, for all ORFs longer than 20 codons a MTS was calculated with MitoFates software. For potential isoforms of annotated proteins (novel or annotated before) a difference in MTS prediction between annotated start codon and potential aTIS was calculated to assess possibility of initiation dependent MTS translation.

### tRNA Copy Numbers and Heptamer Frequency Analysis

tRNA copy numbers for the reference genome (DMKU3-1042) were determined with tRNA scan-SE (Chan and Lowe, 2019). The following formula was used for all heptamers found in CDS regions, where B is the +1 nucleotide (7^th^ base of heptamer).

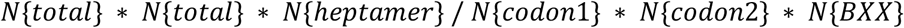

## RESULTS

### Generation of Multiomic Data

We wanted to generate a holistic view of gene expression in *K. marxianus* to understand the coding potential of the genome and to explore its potential for variation. To do this, we employed an integrated combination of transcriptomics and ribosome profiling in what can be called a “multi-omics” analysis since a range of different techniques are used to analyse the resulting data (Figure 1). Transcriptome analysis was performed with now-standard RNA-Seq methods and for ribosome profiling in *K. marxianus,* we applied a custom protocol that is a modified version of that used in *S. cerevisiae* (McGlincy and Ingolia, 2017; Fenton *et al.*, 2022). To try to capture expression of as many genes as possible, our experimental design involved the collection of ribosome profiling and RNA-seq data from cultures grown at 30°C, at 40°C. and at 5, 15, 30 and 60 minutes after a transfer from 30°C to 40°C. Using the updated gene annotation that is described later, we detected the translation of 4872 protein-coding genes with at least 20 mapped ribosome footprints (RPFs), representing ~95% of the previously annotated protein-coding genes in *K. marxianus*. Ribosome profiling data displayed excellent triplet periodicity, allowing us to interpret which frame was translated in a particular locus (Figure 1).

**Figure 1.**
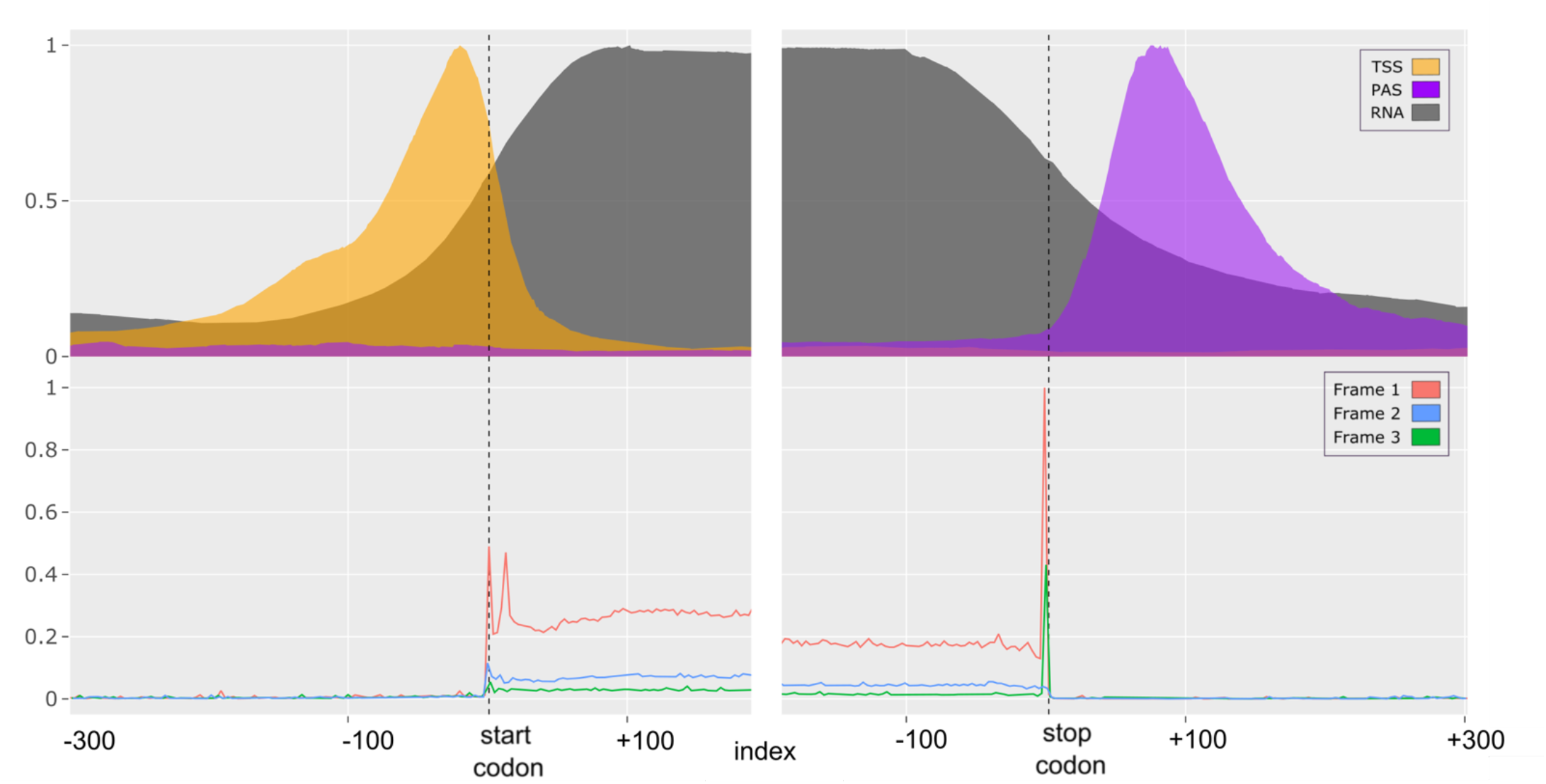
Multiomics metagene profile plot for all annotated protein coding genes relative to start and stop codons. The top plot (transcriptomic data) shows densities of reads from TSS (transcript start sites), PAS (polyadenylation sites) and RNA-seq experiments. The bottom plot shows densities of ribosome footprints differentially coloured based on the best supported reading frame with Frame 1 corresponding to the frame of annotated CDS. Vertical dashed lines represent the start and stop codons of CDS regions. Outward CDS boundaries include 300 nt from the start or stop codon. Inward CDS boundaries extend 200 nt downstream of CDS start and 200 nt upstream of CDS stop.

While ribosome profiling identified the regions of genes that were translated, we also wanted to map the 5’ and 3’ ends of mRNAs. Since the RNA-seq reads generated in our experiments are not suitable for accurate mapping of transcription start sites (TSS), we made use of TSS-seq data that was available from a previous study with the strain DMKU3-1042 (Lertwattanasakul *et al.*, 2015). From the published data, it was possible to precisely map the transcription start sites (TSS) for 2901 *K. marxianus* genes (57% of the total). The majority of genes displayed a single TSS but 489 genes had two TSS and ~150 had three or more TSS (Supplementary Figure 1). While this level of heterogeneity is comparable with that seen in *S. cerevisiae* (Arribere and Gilbert, 2013), it was found that the median 5’ leader length of 97 nt was almost twice that reported in *S. cerevisiae* (Nagalakshmi *et al.*, 2008). In some cases (407 genes), TSS were detected within the coding sequence indicating that transcriptional variation can give rise to alternative shorter mRNA isoforms. To locate polyadenylation sites (PAS) on the 3’ends of mRNAs, we designed a computational approach to identify and separately align RNA-seq reads with polyA tags (using our polyA-enriched RNA-seq data, which was suitably 3’ biased for this purpose). We mapped polyadenylation sites (PAS) for 4682 genes (91% of the total), of which ~3000 genes contain a single PAS and ~1700 genes had more than one PAS (Supplementary Figure 1). The median distance between the stop codon and the PAS was 128 nt, similar to values reported in *S. cerevisiae* of 104 nt (Nagalakshmi *et al.*, 2008) or 166 nt (Ozsolak *et al.*, 2010), indicating that the 3’ trailer lengths are comparable in both species. As with TSS, PAS mapped within the CDS for some genes. Indeed, 607 genes have an internal PAS, and for 335 of these genes it is a major site for polyadenylation. The presence of 5’ truncated transcript isoforms have been reported previously (Arribere and Gilbert, 2013), but the exact roles of these transcripts remain elusive and the same study discovered many of these truncated mRNAs are subject to nonsense mediated decay (NMD) due to out of frame initiation and termination at premature stop codons. Together, these data revealed the genome-wide locations of transcript start sites and polyadenylation sites (Supplementary tables 1 & 2), allowing us to accurately determine the boundaries of expressed mRNAs in *K. marxianus*.

### Multiomics analysis reveals potential gene regulation and expression of alternative proteoforms

Combining ribosome profiling data with accurate transcript start and polyadenylation site positions, we investigated the extent to which variability in transcription, polyadenylation and translation are responsible for the synthesis of alternative proteoforms in *K. marxianus*. We created a computational pipeline that uses ribosome profiling data to call all potential translated ORFs outside of canonical annotated protein coding genes, and then classified them depending on their position relative to known protein coding genes. Parameters such as P-site scores, ORF length, distance to parent gene and number of RPFs were used to filter the data (see Methods and Supplementary Figure 4). This pipeline identifies features such as N-terminal extensions (NTE), internal ORFs (iORF), upstream open reading frames (uORF), overlapping upstream open reading frames (ouORF), antisense translation (aORF) and ribosome frameshifting (see Supplementary Figure 5 for visualization of these ORFs). These were systematically explored on a genome-wide level and in each candidate case, manual multiomics visualization around each locus of interest confirmed the presence of the reported feature. Full lists of genes displaying these features are provided in Supplementary tables 3 – 7 and specific examples of each are described below.

#### 1. Expression of alternative proteoforms

Proteins with alternative N-terminus (PANTs) can arise due to regulation at the level of transcription or translation. Use of an alternative transcription start site (aTSS) can give rise to a longer or a shorter mRNA isoform and the consequential use of a different translation start codon. In contrast, the use of alternative translation initiation sites (aTIS) arises because of recognition of different start codons, for example *via* leaky scanning. Both types of PANTs were evident in *K. marxianus* and we detected a total 5499 potential PANTs in 4825 genes (Supplementary table 3). It is known that N-terminally extended (NTE) proteoforms are sometimes used in *S. cerevisiae* to encode mitochondrial targeting signals (MTS) that localise to the mitochondrion (Monteuuis *et al.*, 2019), therefore we used mitochondrial signal prediction methods to assess our NTE candidates (see methods). This analysis predicted that 314 NTEs may encode a MTS (Supplementary table 4).

Several examples of *K. marxianus* genes that use either aTSS or aTIS to generate PANTs are depicted using multi-omics plots in Figure 2. In the case of three genes (*FUM1*, *FOL1* and *TRZ1*) the PANTs control mitochondrial localisation, and for the fourth (*ADO1*), the function of the alternative isoforms is not known. *FUM1*, encodes fumarase, which can be located in either the cytoplasm or mitochondrion in *S. cerevisiae* (Wu and Tzagoloff, 1987). In *K. marxianus*, two distinct TSS are visible (orange peaks), one of which overlaps the annotated start codon (Figure 2A). There is clear evidence of translation from the annotated start codon, however this is most likely to occur from the longer mRNA isoform as the position of the downstream TSS precludes use of this AUG codon for initiating ribosomes assembling at the 5’end of mRNA. There is a second in-frame AUG codon 51 nts downstream of the first, and the large increase in RPFs downstream of this AUG codon is a strong indication that translation initiates here. While it is possible that this second AUG codon is accessed *via* leaky scanning, it is not likely because the first AUG codon is in a strong Kozak context, and thus, the shorter proteoform is probably translated from an mRNA isoform made using the downstream aTSS. The peptide sequence between these two codons is predicted to be a mitochondrial targeting signal (MTS) and the data suggest that the localisation of Fum1p to the mitochondrion or the cytoplasm is regulated at the level of transcription. This differs from *S. cerevisiae*, where it has been proposed that regulation of protein folding *via* intracellular metabolites determines localisation of Fum1 (Herrmann, 2009; Regev-Rudzki *et al.*, 2009). In contrast to Fum1, Fol1, encoding a multifunctional enzyme involved in folic acid biosynthesis, is an example of a protein where PANTs with, or without, an MTS arise due to the use of alternative translation start sites (aTIS) on a single transcript (Figure 2B). In this case, there is a single TSS but clear evidence of a low amount of translation initiation upstream of the annotated AUG. This translation initiates from a near-cognate UUG codon, which is likely to be inefficiently recognised, causing leaky scanning and subsequent initiation at the downstream AUG. Further confirmatory analysis of the use of these two translation start sites is presented in Supplementary Figure 6. Interestingly, Fol1 was reported to be exclusively located in the mitochondrion in *S. cerevisiae* (Guldener et al, 2004), again raising questions as to possible differences between *K. marxianus* and *S. cerevisiae*. *TRZ1*, which encodes a tRNA endonuclease that is localized to both the nucleus and mitochondrion in *S.* cerevisiae (Chen *et al.*, 2005; Skowronek *et al.*, 2014), is another example of where PANTs arising from leaky scanning of an non-cognate start codon (Figure 2C). Again, there is a single TSS but in this case, translation starts at a GUG codon 27 codons upstream of the annotated AUG start codon, which allows incorporation of the MTS. It is noteworthy that in *S. cerevisiae*, ribosomes initiate translation at an upstream CUG codon in the *TRZ1* mRNA (Monteuuis *et al.*, 2019). While the localization signals are generally short, we also find examples of PANTs with considerable length variation at their N-termini. An example is *ADO1*, encoding an adenosine kinase, where we found evidence for aTSS giving rise to PANTs differing by 100 AA (Figure 2D). Here, both RNA-seq and Ribo-seq coverage suggests the presence of two translated mRNA isoforms with the shorter isoform being more abundant. Indeed, in the figure, both the first TSS and the translation from an upstream AUG are just visible on this scale. Triplet periodicity supports the premise that the upstream AUG is used. No significant matches were found for the extended region in Conserved Domain Database (CDD), nor among protein families in InterProScan but the transmembrane topology and signal peptide predictor Phobius (Käll, Krogh and Sonnhammer, 2004) detected a signal peptide present at the N-terminus of the longer proteoform Supplementary Figure 7) suggesting that the two proteoforms may have different compartmentalisation. *S. cerevisiae* Ado1 lacks this extended region, but it is present in other *Kluyveromyces* species, indicating that the extended Ado1 variant may have a specific, though as-yet unknown, function in *Kluyveromyces* spp.

**Figure 2.**
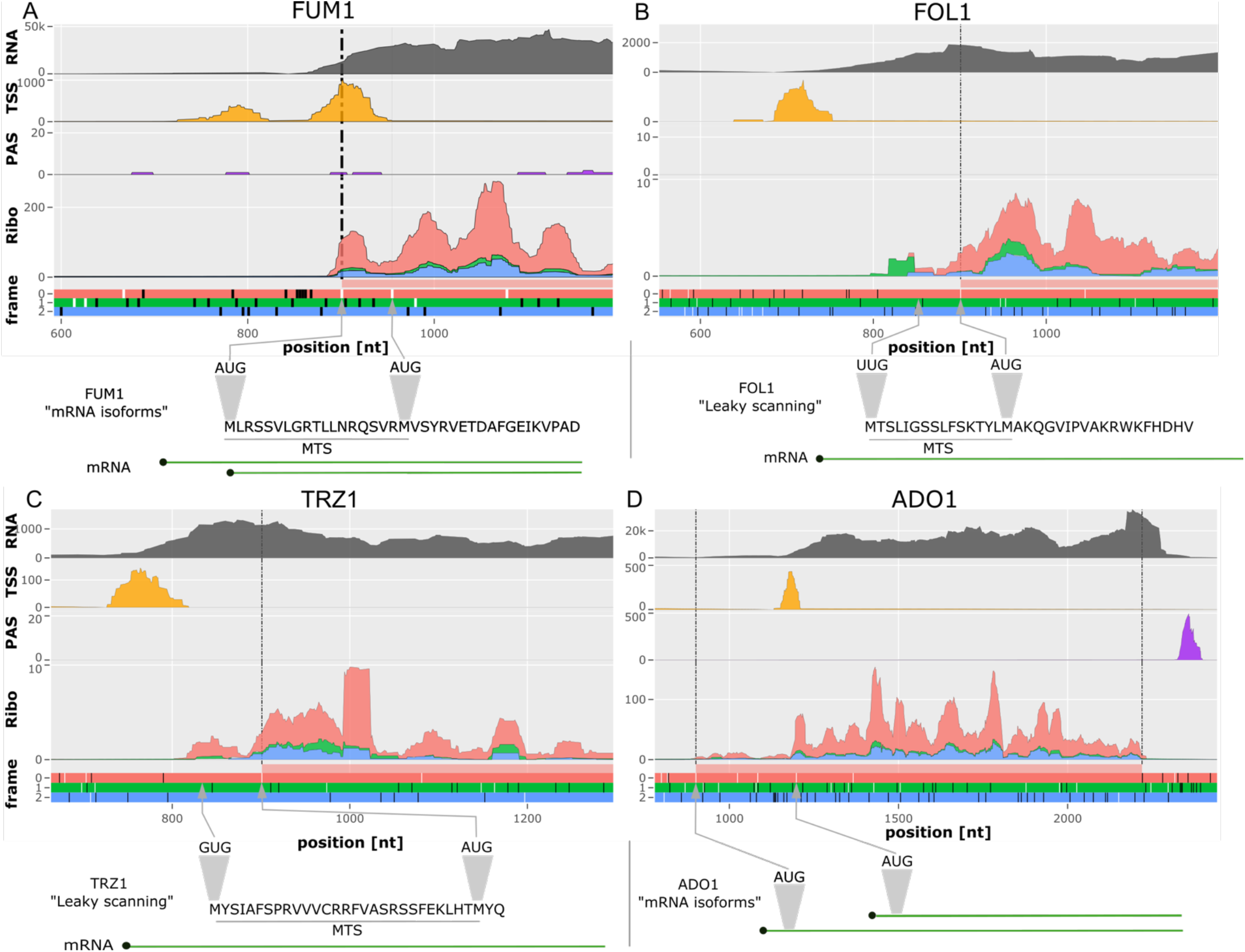
Multiomics evidence of PANTs. Upper panels represent densities of sequencing reads obtained with different techniques, coverage is displayed with the stacked method. For ribo-seq data, RPFs are differentially coloured to match reading frames in the ORF architecture plot below with dotted line showing the positions of the start of annotated CDS (in red). In ORF plots white lines represent AUG codons while black lines represent one of three stop codons. The bottom schematic illustrates suggested models of PANTs expression. A. *FUM1* exemplifies PANTs expression from two RNA isoforms where the location of the most 5’ AUG codons differ. B. *FOL1* exemplifies PANTs expression from the same mRNA where initiation at two different starts (UUG and AUG) occurs due to leaky scanning. A more granulate view of translation initiation from the upstream UUG is shown in supplementary figure 6. C. *TRZ1* exemplifies PANTs expression from GUG and AUG start codons. D. *ADO1* exemplifies PANTs expression via two mRNA isoforms both utilizing AUG start codons. *ADO1* exemplifies PANTs expression via two mRNA isoforms both utilizing AUG start codons.

Translation of internal ORFs (iORFs), which arise from alternative translation of the +1 or −1 frame (relative to the AUG codon) within the annotated CDS of a gene, is another mechanism that cells use to generate alternative proteoforms. We found 861 potential instances of this in *K. marxianus* (Supplementary table 5) and examined several in more detail to illustrate the depth of data that can be retrieved from the multi-omics analysis. *EST3* is an example where such internal translation could be due to leaky scanning past the annotated start codon with initiation at a downstream AUG codon in a different frame (Figure 3A). Initiation at the first AUG would give rise to the functional Est3 protein whereas initiation at the second AUG leads to the translation of an out of frame 24 AA peptide. The poor context (<u>U</u>CC<u>AUG</u>CCC) of the first AUG codon makes it likely that a proportion of scanning ribosomes fail to recognize the main start codon and initiate at the second downstream AUG. An alternative process whereby scanning ribosomes can slide from one AUG to another during the initiation process while awaiting a critical GTP hydrolysis step has been described in mammalian systems that when two AUG start codons are in close proximity, and such a mechanism cannot formally be ruled out here (Terenin *et al.*, 2016), Regardless of the precise mechanism, it is seen that there is weak translation of the first ORF (Figure 3A, red RPFs), which encodes functional Est1 and much stronger translation of the second ORF, encoding the 2AA peptide (Figure 3A, green RPFs) and Interestingly, in *S. cerevisiae* and most other yeasts, *EST1* utilizes +1 frameshifting and it was previously noted that the *Kluyveromyces* genus is an exception that does not utilize frameshifting (Farabaugh *et al.*, 2006). Both +1 frameshifting in *S. cerevisiae* and suboptimal initiation in *K. marxianus* are expected to result in low translation efficiency of the *EST3* mRNA, and thus it appears that different yeast lineages have arrived at distinct mechanisms to translationally restrict the level of Est3. One can speculate that this may be a regulated process whereby some stimulus would act to overcome the translational controls allowing production of higher amounts of the protein.

**Figure 3.**
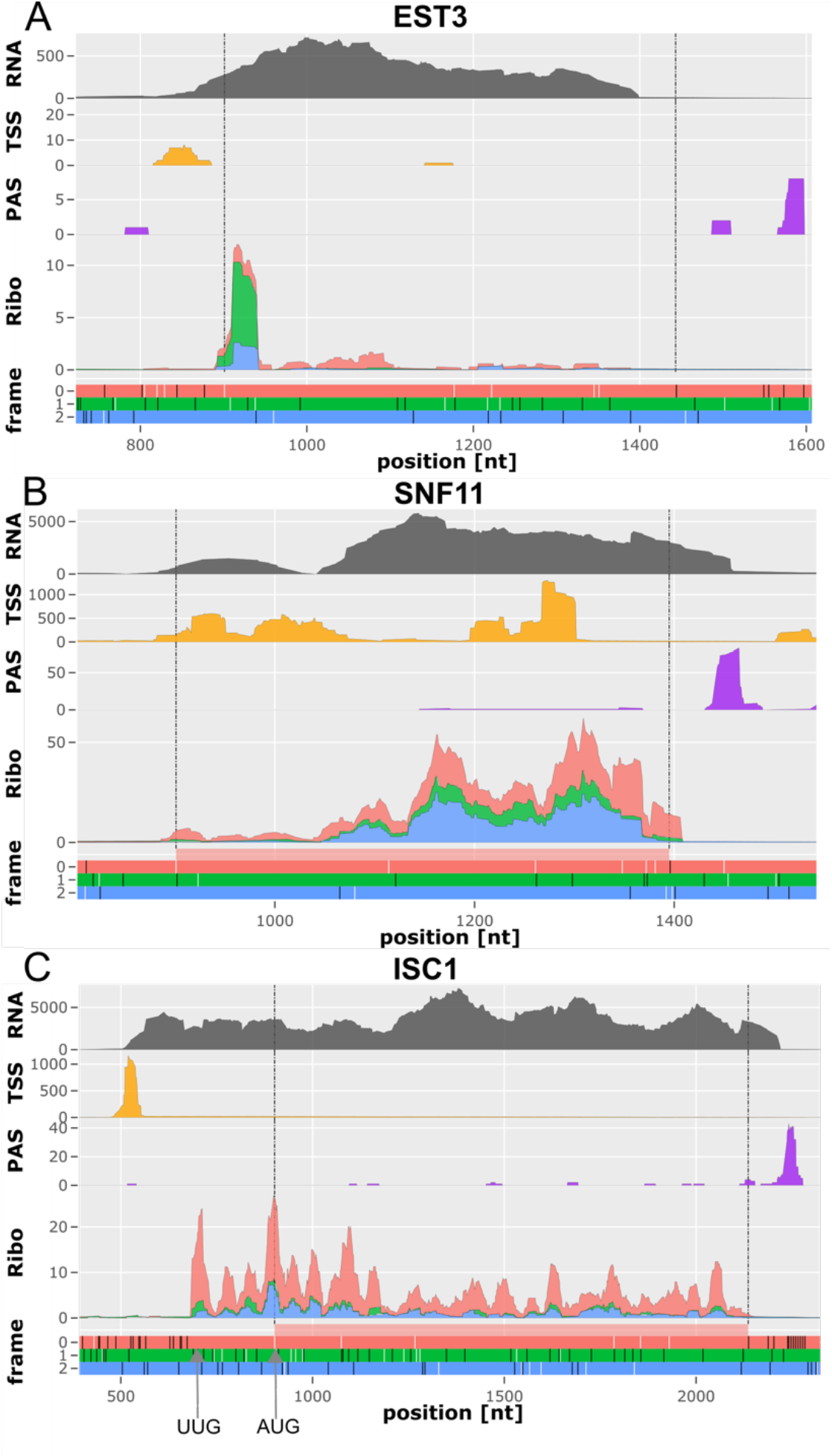
Multiomics plots display out of frame internal translation initiation of known genes (iORFs) and exclusive non-AUG translation initiation. A. *EST3* main CDS is encoded in red frame and described iORF is in green (+1 frame). B. *SNF11* locus is encoded in zero (red) frame while iORF is encoded in the −1 (blue) frame. C. Multiomics plot of the *ISC1* locus. The annotated CDS is in the red frame. The proposed exclusive UUG and annotated AUG start codon are marked below the frame track. See Figure 2 for explanation of the multi-omics plots.

A different type of iORF is seen at the *SNF11* locus, which encodes a subunit of the SWI/SNF chromatin remodelling complex (Figure 3B). In this case, translation from the first (annotated) AUG gives rise to Snf11 (165 AA) but there is also evidence of both additional transcription starts and translation initiation from other AUG codons downstream of the annotated AUG. In fact, the TSS data suggest several sites of transcription initiation that would lead to the production of multiple mRNA isoforms shorter than the annotated one. These isoforms lack the annotated *SNF1* start codon and are likely to be translated from AUG codons further downstream. This idea is supported by the increased ribosome profiling density in the area of the second in-frame AUG (Figure 3A, red RPFs) as well as around a −1 frame AUG codon located a few nucleotides upstream of the second in-frame start codon (Figure 3A, blue RPFs). Use of these alternative AUGs would give rise to N-terminally truncated proteoforms in the case of in-frame codons, and a 92 AA peptide if the −1 frame was used. For the latter, the predicted protein does not have any homologs in databases so its significance remains to be established.

*ISC1*, encoding Inositol phosphosphingolipid phospholipase C, is an intriguing case that reveals unusual use of non-AUG initiation codons in *Kluyveromyces* spp. Initial analysis suggested that this locus might encode a PANT as translation upstream of the annotated AUG start codon was evident (Figure 3C). Deeper analysis, however, revealed that translation occurs exclusively from an upstream UUG start codon, which is in good Kozak context (AAG_UUG_ACG). TSS and RNA-seq confirms that the *ISC1* locus encodes a single major transcript isoform. The possibility that leaky scanning could allow the annotated AUG codon be used is discounted as there are multiple of out-of-frame AUG codons between the TSS and this AUG. We excluded the possibility that the observed initiation at UUG was an artefact due to a recent mutation of the UUG to an AUG in the strain that we used to generate ribosome profiling by examining the sequence of RPFs aligned to this region and confirming that there were no mismatches. To determine if this exclusive non-AUG translation is conserved, we aligned the *ISC1* protein sequences from *K. marxianus*, *Kluyveromyces lactis* and *S. cerevisiae* and also examined the locus architecture (Supplementary Figure 8). The architecture in *K. lactis* matches *K. marxianus* but, in *K. lactis*, the predicted start codon is a GUG. The start codon Kozak context of both GUG (*K. lactis*) and UUG (*K. marxianus*) are identical (AAA_NUG_ACG, where N is G or U) and the N-termini of the proteins are conserved, providing a strong indication that this region is translated in both yeasts. We also used Trips-Viz and ribosome profiling data from multiple published studies at the *S. cerevisiae ISC1* locus to examine translation in this species (Supplementary Figure 8). There is no evidence of the use of non-AUG translation but we did observe that the start codon is incorrectly annotated in *S. cerevisiae*, with in-frame translation starting at an AUG codon downstream of the annotated start codon (see Supplementary Figure 8). For the alignments shown in Supplementary Figure 8, the previously predicted upstream 29 amino acids of the *S. cerevisiae* Isc1 were removed as these are not translated. Despite the difference in the choice of start codon, there is evidence of conservation of the N-termini between *Kluyveromyces* and *Saccharomyces*, raising further questions as to why *Kluyveromyces* has evolved to use non-cognate start codons for this gene (Supplementary Figure 9).

#### 2. Translation of short ORFs within 5’ leaders

Upstream open reading frames (uORFs) are one of the best-studied mechanisms by which translation of an mRNA is regulated in yeast and fungi (Hood *et al.*, 2009). In addition, across eukaryotes, many uORFs are known to regulate translation in response to various stimuli, such as polyamines in mammals (Law *et al.*, 2001; Ivanov *et al.*, 2018; Vindu *et al.*, 2021) and plants (Franceschetti *et al.*, 2001), magnesium levels in mammals (Hardy *et al.*, 2019), boron in plants (Tanaka *et al.*, 2016) and arginine levels in yeast (Gaba, Jacobson and Sachs, 2005) among many others. In our analysis of the *K. marxianus* data, we detected 818 uORFs that do not overlap with CDS (see Supplementary table 6). These included *GCN4,* which is considered a paradigm for translational regulation via uORFS in yeast (Hinnebusch, 2005) where several short uORFS are translated, supporting the idea that the *GCN4* regulatory system in *K. marxianus* is identical to that of *S. cerevisiae* (Supplementary Figure 10). A further example to illustrate this type of uORF is presented in Figure 4A, where strong initiation at an AUG-initiated uORF ~140 nt upstream of the main CDS for *SNG1* is seen. *SNG1*encodes a protein involved in drug resistance so it also fits the pattern of this type of regulation being used for some stress-response genes (García-López et al., 2010). We also identified 443 potential overlapping uORFs (ouORFs), which we define as cases where the uORF overlaps the main ORF/CDS and thus translation is expected to be mutually exclusive (Supplementary table 7). *RAD59*, encoding a DNA repair protein provides an example of this, where an ouORF (Figure 4B, blue RPFs) overlaps the main ORF (Figure 4B, red RPFs).

**Figure 4.**
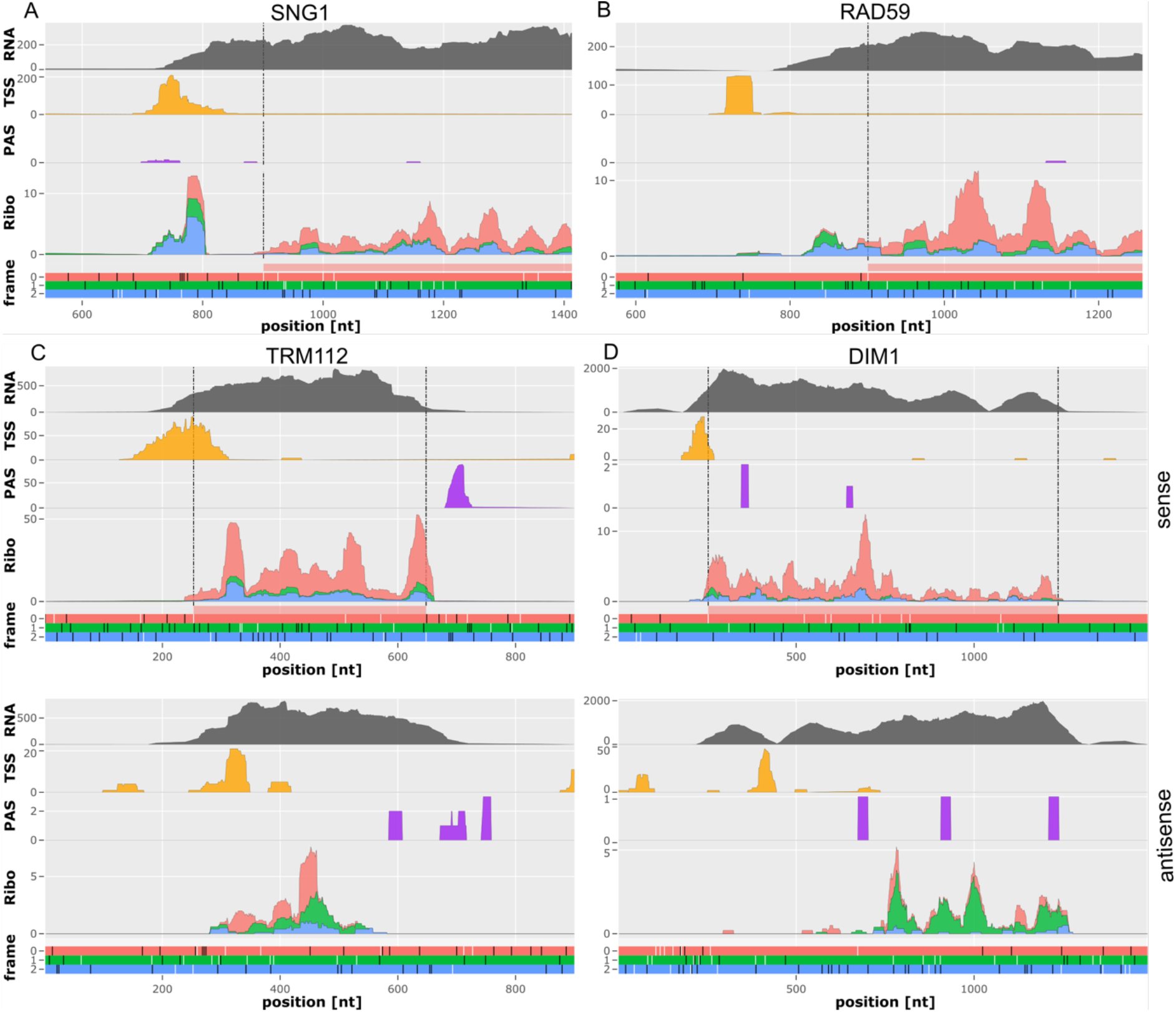
Translation within 5’ leaders and antisense translation of protein-coding genes. A. uORF in the 5’ leader of the *SNG1* mRNA in blue frame (left). B. uoORF in the *RAD59* mRNA in the blue frame which overlaps the main CDS (right). Multiomics plots displaying sense multiomics of C. *TRM112* and D. *DIM1* (top row). Lower row displays exact antisense locus (5’ to 3’) For TRM112, antisense translation can be seen predominantly in two ORFs (red and green) while for DIM1, antisense translation mostly occurs in the green frame. See Figure 2 for explanation of the multi-omics plots.

#### 3. Translation of Antisense mRNA

Antisense translation represents a class of detected translated ORFs (aORFs) on the opposite strand of an annotated protein coding gene. Using GWIPS-viz, we unexpectedly observed that RPFs map to sense and antisense strands of *TRM112* and *DIM1*, suggesting the existence of translated antisense transcripts (Figure 4C and 4D & Supplementary Figure 11). We therefore decided to include aORFs in our computational pipeline to find more candidates for antisense translation. We ranked genes by the number of RPFs aligned to the opposite strand of all protein-coding genes and then validated candidates using multiomics visualisation. This approach uncovered potential antisense expression in 938 genes including *TRS23*, *VPS9*, *bioA* and *EXO84*, which were confirmed with multiomics plots and GWIPS-viz (Supplementary table 8). Antisense translation was previously noted by Duncan and Mata in ribosome profiling data from *S. pombe* (Duncan and Mata, 2014) but its biological significance remains unknown.

#### 4. Novel +1 ribosomal frameshifting at the KLMX_30357 locus

We uncovered a potential +1 frameshifting event during translation of an mRNA derived from the *K. marxianus KLMX_30357* locus (Figure 5A). This locus was previously annotated as two separate protein coding genes (*KLMA_30367* and *KLMA_30368*) but we show that it is a single transcript as there is only evidence for one TSS and 1 PAS. Furthermore, although there was relatively uniform RNA-seq distribution along the length of this putative single transcript, the reading frame changed to the +1 position in the spacer region between the “KLMA_30367” and “KLMA_30368” CDS. This suggested to us that translation of “KLMA_30368” CDS could be due to ribosomal frameshifting. Indeed, sequence analysis revealed the presence of a likely shift-prone pattern (GCG_AGG_C) at the site where the ribosome density in the +1 frame increases (Supplementary figure 12). This particular heptamer sequence was previously shown to support +1 frameshifting in *S. cerevisiae* (Sundararajan *et al.*, 1999) and may be described as a hybrid of Ty1 (CUU_AGG_C) and Ty3 (GCG_AGU_U) +1 frameshift heptamers described in *S. cerevisiae* (Clare, Belcourt and Farabaugh, 1988; Farabaugh, Zhao and Vimaladithan, 1993). Frameshifting prone patterns are known to be underrepresented in coding sequences as they reduce processivity of translation (Shah *et al.*, 2002; Gurvich *et al.*, 2003). We analysed the occurrence of all in-frame heptamers in *K. marxianus* and found that this heptamer occurs far more rarely than what could be predicted based on its codon composition (see methods and Supplementary Figure 12) and is among the ~2% of the rarest heptamers present in coding regions (Supplementary Figure 14). It has been suggested that severe imbalance between availability of tRNAs for the A-site codons in 0 (AGG) and +1 (GGC) frames in Ty1 frameshifting site is responsible for its high efficiency in *S. cerevisiae* (Baranov, Gesteland and Atkins, 2004). Based on the assumption that tRNA copy number correlates with tRNA abundance (Percudani, Pavesi and Ottonello, 1997), we generated a table that estimated the relative abundance of each tRNA in *K. marxianus* (see methods and Supplementary table 9). We find that the ratio of tRNA^Arg^_CCU_ decoding AGG (0 frame) to tRNA^Gly^GCC decoding GGC (+1 frame) is 1:9, which is consistent with a hypothesis that a tRNA imbalance may be the driver of this frameshift. The expected full-length product with +1 frameshifting would be a protein 857 amino acids in length. We searched for homologs of this full length protein and discovered this gene can be separated to three regions of interest (Figure 5B). The zero-frame ORF is a homolog of *S. cerevisiae YLR257W*, a gene of unknown function. The +1 frame product contains a midasin/AAA ATPase domain except for the C-terminus, which is highly similar to the C-terminus of *S. cerevisiae AIP5*. Aip5p is part of a multiprotein polarisome complex which catalyses the formation of actin cables for polarized cell growth during budding (Glomb, Bareis and Johnsson, 2019; Xie *et al.*, 2019). Interestingly, the C-terminus of Aip5p has been shown to be responsible for this activity (Figure 5B). Thus, it is possible that this gene is translated into two proteoforms with distinct functions. We note *S. cerevisiae* Aip5 also contains a midasin/AAA ATPase domain upstream of the C-terminus domain, like the +1 product.

**Figure 5.**
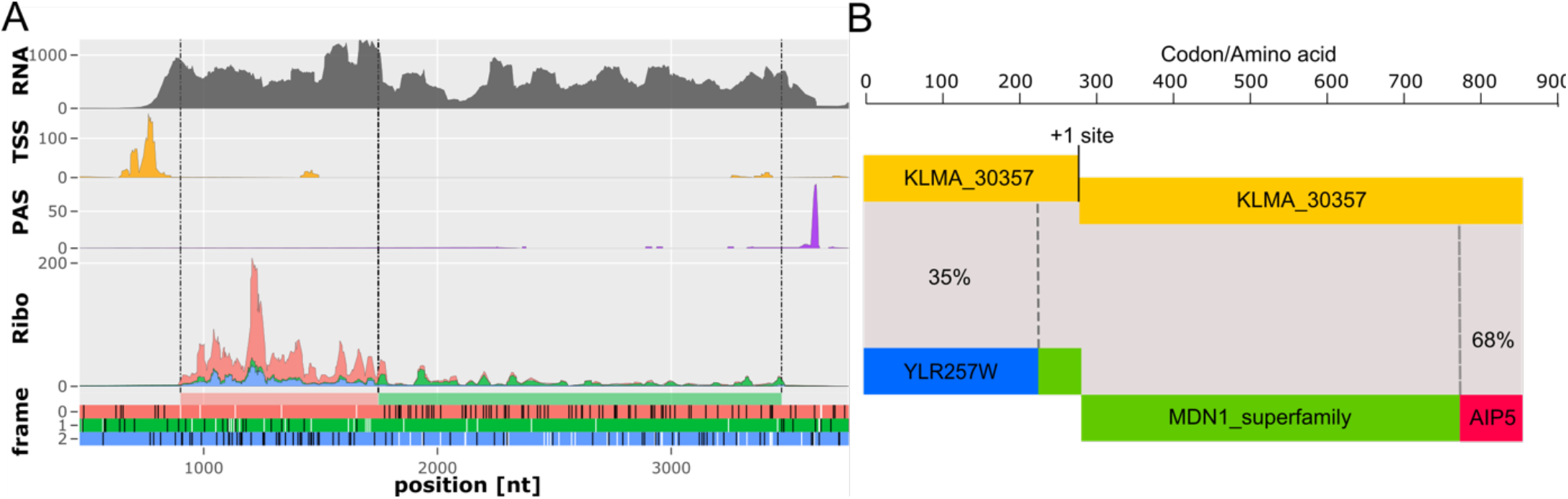
Frameshifting at the KLMX_30357 locus. A displays the region including the newly annotated KLMX_30357 0-frame (red) and +1-frame CDS (green) (originally annotated separately as KLMA_30367 and KLMA_30368). B. Schematic of the KLMX_30357 locus and similarity of full-length frameshift products to known genes and superfamily. Percentages refer to the identity with known homologs in *S. cerevisiae*. See Figure 2 for legend explanation of the multi-omics plots.

### Detection of novel protein coding genes and improvement of existing genome annotation

During analysis of the ribosome profiling data, it was apparent that there were significant numbers of RPFs aligning to the regions of the DMKU3-1042 reference genome that had not been annotated as protein coding. Some of these were found to be caused by a ~40 Kb annotation gap on chromosome 1, which excluded 20 genes, but there were multiple others not explained in this way. Therefore, we decided to use our ribosome profiling data to improve the *K. marxianus* DMKU3-1042 genome annotation by creating a semi-supervised pipeline to detect translated protein coding regions that were not annotated as genes in the reference genome. With this approach, we discovered 171 unannotated candidate genes in the DMKU3-1042 genome and further investigated these putative genes by generating sub-codon ribosome profiles for each transcript using Trips-Viz (Kiniry *et al.*, 2019, 2021). These plots were explored manually for the consistency of ribosome profiling density and triplet periodicity in case artefacts had been introduced by the computational pipeline (see Supplementary Figure 15 for examples). We also generated a table displaying the periodicity score and the number of RPFs per open reading frame for each of these candidate genes (Supplementary table 10). In all cases, the patterns are consistent with protein encoding genes. We then analysed the amino acid sequence to explore this novel gene set. Each of the 171 CDSs was conceptually translated and the resulting protein sequences were queried against the NCBI non-redundant protein database with the BLASTP tool (Altschul *et al.*, 1990). Finally, we interrogated the annotated genomes of 18 yeast species within the budding yeast sub-phylum (*Saccharomycotina*) using both BLASTP and TBLASTN (Figure 6). BLASTP can be used to identify homologs in existing annotations, while TBLASTN allows for identification of homologs in the corresponding genomes irrespective of the completeness of its annotation. Thus, an existence of a high scoring hit in a TBLASTN search that is absent in a BLASTP search would indicate the presence of an ortholog that has not been annotated. This analysis included species in the *Kluyveromyces* genus, other species in the *Kluyveromyces / Lachancea / Eremothecium* (KLE) clade, representatives from the *Zygosaccharomyces / Torulaspora* (Z/T) clade, and three post whole-genome duplication (WGD) species, *Candida glabrata*, *Kazachstania africana* and *S. cerevisiae* (Shen *et al.*, 2018). A larger number of *Lachancea* species were included as this is the most closely related genus to *Kluyveromyces*. There were substantial differences between the results of the BLASTP and TBLASTN analyses, indicating that quite a number of protein coding genes were missing from the genome annotations of some species. For the majority of the 171 genes, we found homologous proteins in most or all of the 17 other yeast species from the budding yeast subphylum that we included. In some cases, these genes encoded orthologs of proteins with known or easily predicted functions; for example, transcription factors, ribosomal proteins, a sugar transporter, a redox protein, heat shock proteins and others (Supplementary table 11). The addition of these protein coding genes increased the total number of annotated protein coding genes from 4,952 to 5,118 in the DMKU3-1042 genome (see Supplementary table 12 for comparison of published *K. marxianus* genomes). Ultimately, of the 171 newly identified translated genes, only 10 genes are specific to *K. marxianus* and a further 16 genes appear specific to the *Kluyveromyces* genus. Analysis of these 26 proteins with the PFAM superfamily database failed to identify any known domains that would give clues as to function.

**Figure 6.**
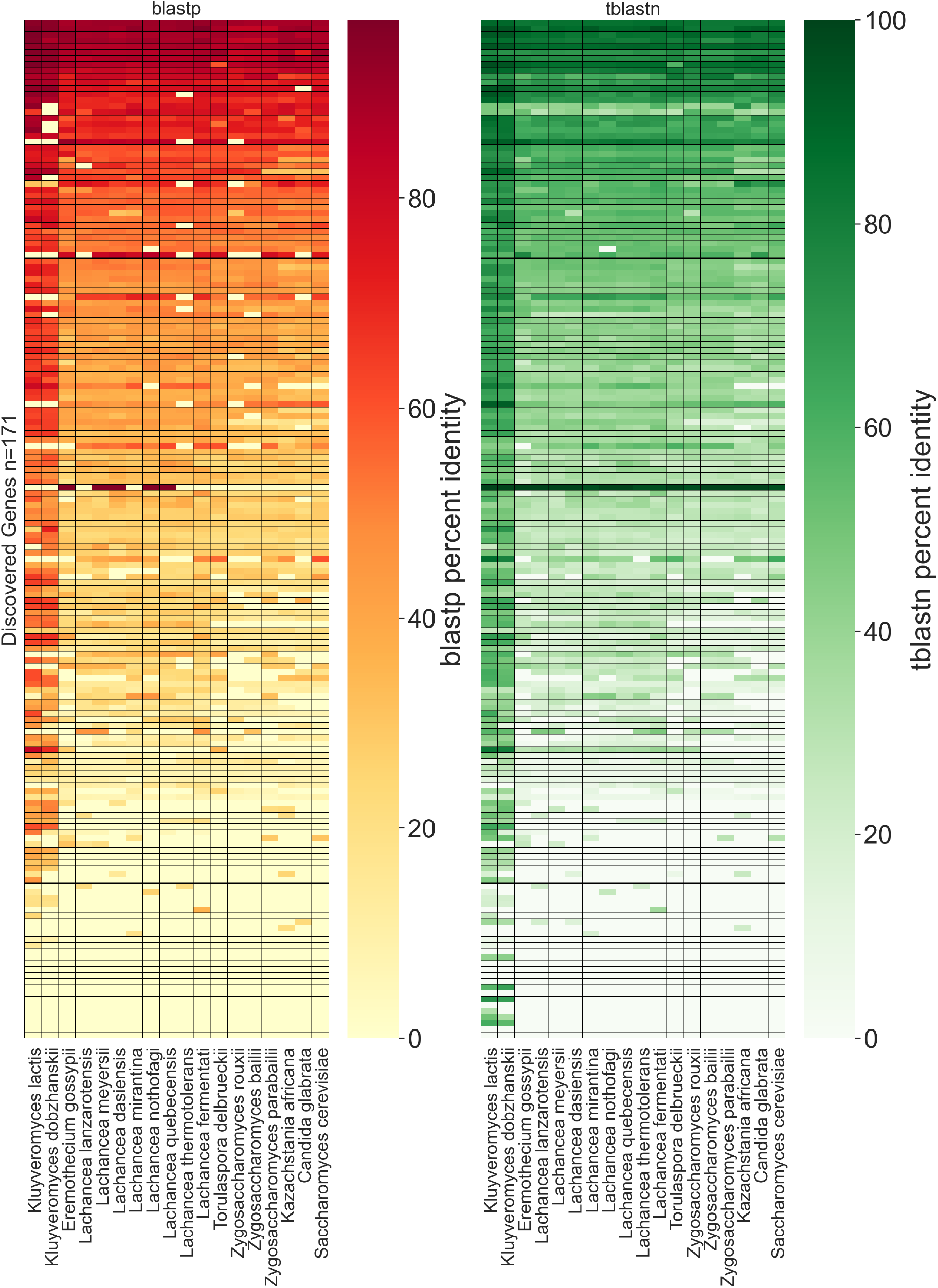
Analysis of orthologous groups for 171 newly identified genes. Left is a heatmap of % amino acid identity for the BLASTP hits obtaining for a search against proteins annotated in other genomes. Right is a heatmap of % amino acid identity for the TBLASTN hits obtained for a search against genomic sequences. Each row in both tables corresponds to the same gene for comparison.

In addition to discovering previously non-annotated genes, we found and corrected 120 incorrectly annotated genes. A full list and breakdown of these gene corrections is provided in Supplementary table 13. The start codon was reannotated for 69 genes as an unequivocal density of in-frame translating ribosomes could be observed starting either upstream or downstream of the previously annotated start codon (for example see Supplementary Figure 16 & Supplementary table 14 for description of all start codon corrections). Multiple corrections were also made in nuclear-encoded intron-containing genes. The total number of annotated spliced genes increased by 13 to 183, as 28 of the newly-annotated genes contain introns but 15 genes previously annotated as containing introns did not, in fact, contain introns. In most of these cases, translation initiated from an in-frame AUG in the reported “second” exon and there was no evidence of the “first” exon being translated. One example of a spliced gene where the splicing was missed is *QCR9*, which encodes a subunit of Complex III in the mitochondrion electron transport chain (Supplementary Figure 17). In 20 cases where splicing was reported, the coordinates of intron-exon boundaries were incorrect and needed to be amended, for example, the *NUP60* locus (Supplementary Figure 18). A full list of spliced genes in *K. marxianus* is provided in Supplementary table 15. The genes encoding the ribosomal proteins Rpl7, Rps9 and Rpl36B were annotated on the wrong strand and this was corrected (see Supplementary Figure 19 for example). Finally, we correctly annotated the +1 frameshifting genes *OAZ1* and *ABP140* to include both the 0 frame and +1 frame as original annotations reported a single ORF (for example see Supplementary Figure 20), and we also included as a correction the −1 frameshift gene *KAT1* (Rajaei *et al.*, 2014), originally annotated as two separate protein-coding genes. We deposited our updated annotation to our developed RiboSeq.Org resources and users may now freely explore our ribosome profiling and transcriptomic data on the GWIPS-viz genome browser relative to the original or updated genome annotation. In addition, TSS and PolyA coverage tracks have also been added to GWIPS-viz (https://gwips.ucc.ie/).

## DISCUSSION

A robust accurately annotated genome is an important tool in modern yeast genetics and while advances in sequencing technology make it easy to generate a genome sequence, databases are replete with incomplete information and annotation errors. This is particularly the case when considering non-model yeasts since annotation is generally based on using *S. cerevisiae* as the reference. This can lead to an accumulation of errors because of the underlying assumption that the new genome will not deviate substantially from the supposed reference genome. This point is illustrated by a recent study of methods for comparative genomics that concluded that variable annotation results in large-scale over-estimation of lineage-specific genes (Weisman et al. 2022). We demonstrate how a genome annotation can be improved by including experimental data that measure transcription and translation. This multi-omics approach enabled us to significantly improve the *K. marxianus* annotation through the addition of genes, correction of splicing events, and identification of correct transcription and translation start sites. After *S. cerevisiae*, the *K. marxianus* genome annotation presented here is the most accurate and complete genome annotation within the budding yeasts. This annotation can serve as a reference for the annotation of other *K. marxianus* strains and closely related species and the methodology could be applied to other non-traditional yeasts. It may also be valuable for a number of previous studies in *K. marxianus* that used RNA-Seq to study responses to various stresses and stimuli. Over 300 new or corrected gene annotations are now present and, given that species-specific genes are predicted to be important for stress response and niche adaptation, it may be worthwhile to reanalyse RNA-Seq data using this new annotation. Similarly, for comparative genomic studies, proper gene level annotation is crucial to avoid errors and thus this new reference will be valuable.

A major innovation of this study was the integration of different omic methods. The development of a method for ribosome profiling in *K. marxianus* is recent (Fenton *et al.*, 2022) and this work marked its first use. In combination with TSS data and 3’ biased RNA-seq (due to poly(A) selection), it was possible to reveal mRNA boundaries and the presence of mRNA isoforms on a genome-wide scale. This provided information on both the transcriptional and translational landscape, which aided in deciphering events such as N-terminal extensions and uORFs through the use of multiomics visualization of a locus of interest. Our computational pipeline showed the diversity in the proteome with multiomics data, identifying the expression of alternative proteoforms and the translation of many short ORFs. With the annotation of mRNA boundaries, we were able to decipher whether alternative proteoforms are generated by alternative transcription due to the use of different transcription start sites within the promoter (such as with *ADO1* and *FUM1*) or alternative translation initiation due to leaky scanning (*FOL1*).

It will be very interesting in the future to compare the complex and varied genomic diversity in *K. marxianus* with that of *S. cerevisiae* as they represent two yeast lineages with a very different evolutionary history. One, *S. cerevisiae*, arose from a hybridisation between parents from the KLE and ZT clades (Wolfe and Shields, 1997; Marcet-Houben and Gabaldón, 2015), and thus has a duplicated genome with numerous ohnologs. In contrast, *K. marxianus* represents pre-WGD yeasts and thus may be more representative in many regards of the ancestral state. Already from our limited analysis, we identified multiple examples where the mechanism used to generate proteoforms, or to regulate levels of proteins, is different between the two yeast species. In some cases, the *S. cerevisiae* data may be incomplete and some previous assumptions incorrect, but in others, e.g for regulating *EST3*, it is clear that different mechanisms are used. Many hypotheses can be generated from our dataset and are likely to be a fertile area of study to understand evolution and niche adaption in the Saccharomycetales.

Despite the wealth of data and novel insights generated in this work, it must be acknowledged that these data were obtained from a relatively small number of growth conditions. Thus, although we managed to detect expression of the vast majority of protein coding genes, it is possible that some aspects of regulation were missed because they only arise in some specific condition that we did not use. This is, of course, the case with all previous genome-wide studies of gene expression in other yeasts as well. The future application of integrated omics techniques for mapping the ends of transcripts and positions of ribosomes under different conditions can be expected to reveal additional features of protein-coding organisation in the *K. marxianus* genome.

## MATERIALS AND METHODS

### RNA-Seq and Ribosome Profiling

RNA-Seq and ribosome profiling was carried out as in (Fenton *et al.*, 2022). Splice junctions were identified for novel intron containing genes and genes which required splice site corrections using the splice aware STAR RNA-seq aligner (Dobin *et al.*, 2013). In the ribosome profiling analysis, RPFs that failed to align to the original CDS regions of DMKU3-1042 annotation were aligned to the reference genome. These alignments were then split into windows using Bedtools (Quinlan and Hall, 2010). Windows were ranked based on the number of alignments and the top candidates were visually assessed using a genome browser (GWIPS-viz (Michel *et al.*, 2014)) where we created a database for *K. marxianus* DMKU-1042 genome. For the BLASTP and TBLASTN heatmaps, custom databases containing annotated protein sequences and genome assemblies were created for use with BLASTP and TBLASTN, respectively (Altschul *et al.*, 1990). For BLASTP and TBLASTN, the following parameters were specified, -seg no, -threshold 11, -max_hsps 1 and -outfmt 6. An e value filter of =<0.01 was applied to blast results.

### TSS-Seq

In order to precisely characterise Transcription Start Sites, publicly available TSS-seq data originating from *K. marxianus* DMKU3-1042 was used (Lertwattanasakul *et al.*, 2015). Data were downloaded from the NCBI SRA repository, adaptor removal and quality trimming was performed with cutadapt, followed with rRNA removal and genome alignment with bowtie. While doing this analysis, we noticed that accession numbers of raw data deposited in SRA do not match expression profiles of conditions discussed in the original paper (Lertwattanasakul *et al.*, 2015) and were evidently mislabelled during sequence deposition. For our analysis, we reassigned the samples to the correct condition, detailed in Supplementary material. Resulting alignments were used for transcriptional units (Transcription Start Region -– TSR) detection by clustering reads with Bioconductor package CAGEr (Haberle *et al.*, 2015). TSRs were subsequently assigned to the nearest coding sequence (CDS) with a minimum relative expression cut-off of 0.05 and 1 TPM was applied to filter out lowly expressed TSRs or unreliable clusters (see Supplementary Figure 5).

### Identification of PAS

For identification of polyadenylation sites, we used our own polyA-enriched RNA-seq data (Fenton *et al.*, 2022). Reads that aligned to the genome were discarded as these are sequences that do not contain polyA tails. From the remaining reads, all trailing A nucleotides were trimmed from 3’ ends of reads and aligned once again to the genome revealing PAS. Aligned reads were processed analogously to the TSS-seq reads: clustering has been performed with CAGEr and followed with assignment to nearest CDS with minimum relative expression cut-off of 0.05 and 1 TPM. Remaining clusters were assumed as *bona-fide* polyadenylation sites (PAS, see Supplementary Figure 6).

### Multiomics plots

Briefly, BAM files were processed with ORFik bioconductor package to generate P-site ribo-seq profiles and RNA-seq coverage profiles (Tjeldnes *et al.*, 2021). In the process of mining the data, RiboCrypt: R package NGS data visualization tool was developed. It takes use of ORFik data management and processing and ggplot2 combined with plotly for data display. RiboCrypt GitHub repository is available at https://github.com/m-swirski/RiboCrypt. The profiles are characterised by very sharp peaks, making profiles less-readable when zoomed-out. Thus a sliding window mean of a profile was used to decrease resolution and increase clarity of a picture. To display coverage we employed the stacked method to avoid blurring the plot by area overlapping.

### Non-canonical translation detection

All possible ORFs starting from any one of the near-cognate codons (differing from AUG by one nucleotide, including AUG) in the *K. marxianus* genome and transcriptome were found with ORFik package (Tjeldnes *et al.*, 2021). Naturally, it resulted in finding multiple ORFs sharing a stop codon. Subsequently, a P-site profile was generated for the longest ORF for each stop codon and a set of parameters was calculated for all nested ORFs. P-site score: % of in-frame reads, read count – given as reads per kilobase (RPK), in-frame coverage fold-change between a region between Nth and Nth+1 start codon and first to Nth-1 start codon. Unique P-site score: P-site score calculated for a region between Nth and Nth+1 start codon. Additionally, for all ORFs longer than 20 codons a MTS was calculated with MitoFates software. For potential isoforms of annotated proteins (novel or annotated before) a difference in MTS prediction between annotated start codon and potential aTIS was calculated to assess possibility of initiation dependent MTS translation.

### tRNA Copy Numbers and Heptamer Frequency Analysis

tRNA copy numbers for the reference genome (DMKU3-1042) were determined with tRNA scan-SE (Chan and Lowe, 2019). The following formula was used for all heptamers found in CDS regions, where B is the +1 nucleotide (7^th^ base of heptamer).

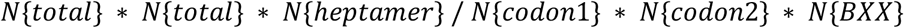

## Data Availability

All Supplementary tables are available at https://zenodo.org/record/6515070#.YnFxEPPMJMM. Ribosome profiling, TSS, PAS and RNA-seq data have been deposited to GWIPS-viz https://gwips.ucc.ie, as well as the original and new annotation tracks. Ribosome profiling and RNA-seq have been deposited to Trips-Viz https://trips.ucc.ie/. Ribosome profiling and RNA-seq datasets have been deposited to the Sequence Read Archive under the under the project accession number PRJEB45612. Our updated genome annotation of *K. marxianus* DMKU-1042 and other relevant files are available at https://doi.org/10.5281/zenodo.6378617.

## ACKNOWLEDGEMENTS

We greatly appreciate the help of Dr Martina Yordanova and Dr Gary Loughran for technical support and advice on developing ribosome profiling.

## AUTHOR CONTRIBUTIONS

P.V.B and J.P.M conceived the study and supervised all aspects of the project. D.A.F performed all wet-lab experiments. M.S integrated *K. marxianus* into RiboCrypt and developed multiomics plots as part of RiboCrypt. D.A.F, M.S and P.O’C carried out bioinformatic analyses. D.A.F and A.M.M developed a database for GWIPS-viz. S.J.K developed a *K. marxianus* database for Trips-Viz. D.A.F, P.V.B and J.P.M prepared the manuscript.

